# Genomic dimensions of Su(H)-targeted regulatory belts in *Drosophila*

**DOI:** 10.1101/055707

**Authors:** Elizabeth Stroebele, Timothy Fuqua, Madelyn Warren, Danielle Herrig, Christian Noblett, Xin Yuan, Albert Erives

## Abstract

Asymmetric Notch signaling promotes divergent fates in select cells throughout metazoan development. In the receiving cell, signaling results in cleavage of the Notch intracellular domain and its import into the nucleus, where it binds Suppressor of Hairless [Su(H)] to promote gene expression in conjunction with contextual cues in the surrounding DNA sequence. To investigate the nature of this contextual logic, we identify 1344 Su(H)-site containing regulatory belts that are conserved across the *Drosophila* genus. Each Su(H)-type regulatory belt (SUH-RB) is a 0.6-1.0 kb chain of conservation peaks consistent with a transcriptional enhancer or core promoter. These regulatory belts contain one or more canonical binding sites for Su(H) along with ~15-30 other binding sites. SUH-RBs are densely clustered in certain chromosomal regions such as the E(spl)-complex, the *Wnt* gene complex, and genes encoding Notch receptor ligands (*Delta and Serrate*). SUH-RBs overlap most known Su(H)/Notch-target enhancers and others, including non-embryonic enhancers that are not identified by embryonic ChIP-seq peaks. Thus, SUH-RBs overcome the stage-specific nature of embryonic ChIP-seq peaks and suggest a pervasive role for contextual tissue-specific pioneer and/or enhancer-licensing factors. SUH-RBs also delineate false positive ChIP-seq peaks, which do not overlap SUH-RBs, are missing even the weakest Su(H)-binding sequences, and have the shortest ChIP peak widths. Last, we characterize several novel enhancers including Su(H)-dependent enhancers at *Notch* and *Delta*, intestinal enhancers at *A2bp1* and *hedgehog*, and distinct enhancers at *roughest*, *E2f1*, and *escargot*.

## INTRODUCTION

The Notch signaling pathway is a key animal innovation central to many developmental operations including: (*i*) tissue compartment boundaries and organizers (signaling across a linear border defining two domains in an epithelium), (*ii*) proneural clusters (signaling across a circular border encircling a field of cells in an epithelium), (*iii*) somatic germline niches (signaling between germline stem cells and neighboring somatic cells), (*iv*) neural stem progenitors and sensory organ precursors (signaling between one cell and surrounding epithelial cells), and (*v*) asymmetric cell fate lineages (signaling between two dividing daughter cells) (Fortini and Artavanis-Tsakonas 1994; Schroeter *et al.* 1998; Voas and Rebay 2004; Ward *et al.* 2006; Liu and Posakony 2012; Housden *et al.* 2014). These operations involve cell-cell signaling between membrane-bound Notch receptor and membrane-bound Notch receptor ligands, such as Delta and Serrate/Jagged. This signaling leads to cleavage of the Notch intracellular domain (NICD) and its import to the nucleus. Nuclear NICD then binds to the transcription factor (TF) Suppressor of Hairless, Su(H), to induce or permit activation of target genes in an amazing array of developmental expression patterns.

Notch signaling to a tissue-specific transcriptional enhancer is frequently integrated with different, context-specific, signaling cues throughout development (Voas and Rebay 2004; Ward *et al.* 2006; Liu and Posakony 2012; Housden *et al.* 2014). These context-specific cues likely include compartment-licensing and pioneer factors, short-range repressors and (longer-range) silencers, as well as other morphogenetic signals coupled to the Notch signal (Stroebele and Erives 2016). Most of these contextual signals are also likely encoded within the enhancer′s DNA sequence, which often encompasses several peaks of conservation consistent with multiple TF binding sites. Insights into contextual modulation of the Notch-signal were uncovered in studies investigating the non-homologous neurogenic ectoderm enhancers (NEEs) at *ventral nervous system defective (vnd), brinker (brk), short gastrulation (sog), rhomboid (rho),* and *vein (vn)* (Erives and Levine 2004; Crocker *et al.* 2008, 2010; Crocker and Erives 2013; Brittain *et al.* 2014). These Notch-permissive enhancers are driven by the Dorsal-Twist dorsal-ventral patterning system, SUMOylation machinery, and the Zelda pioneer factor (Bhaskar *et al.* 2002; Erives and Levine 2004; Markstein *et al.* 2004; Ratnaparkhi *et al.* 2008; Crocker *et al.* 2010; Brittain *et al.* 2014). Thus, the non-homologous NEEs provide an ideal training data set for the development of bioinformatics-methods to identify mechanistically-equivalent sets of Notch/Su(H)-target enhancers, which employ similar contextual logic. Having identified such Notch-target enhancer sets, we wish to determine: (*i*) how Notch is integrated logically with other patterning signals, (*ii*) how developmental signals are specifically acted upon in select tissue compartments despite the frequent use of these signals throughout development, and (*iii*) whether a common theme unifies Notch-regulation across diverse developmental contexts.

Here, we present the computational identification of 1344 Su(H) site containing regulatory belts (SUH-RBs), which are conserved across *Drosophila*. These regulatory belts overlap most known Notch target enhancers and many others not characterized as regulated by Notch signaling and/or Su(H). From this set, we also identify several novel enhancers with distinct site compositions and distinct tissue-specific activities. For example, we characterize novel Su(H)-dependent wing imaginal disc enhancers from the *Notch* and *Delta* loci. We also find that the loci encoding Notch receptor ligands, *Delta* and *Serrate*, correspond to genomic regions with the highest density of SUH-RBs. In contrast, only a single SUH-RB is found at the *Notch* locus, suggesting Notch signaling is heavily modulated by auto-regulatory control of genes encoding Notch ligands. Last, we analyze how the SUH-RBs correspond to stage-specific ChIP-seq peaks, transposable elements, or known *cis*-regulatory modules (CRMs), and find that the SUH-RBs are predominantly composed of unique, stage-and tissue-specific enhancers.

## MATERIALS AND METHODS

### Bioinformatics

**Regulatory belts:** We created a gene regulatory version of the *D. melanogaster* genome assembly (″I-CONTRAST″) by taking the complementary or inverse portion of the intersection of the FlyBase annotated gene set (release 5.51) and the CONTRAST de novo gene prediction data set (see File S1). This intersection was chosen because it was found to result in a genomic track file marking only protein-coding sequences. For example, this filtering step maintains the presence of 5′ and 3′ untranslated regions in the non-protein-coding (regulatory) portion of the genome. We then searched the *D. melanogaster* I-CONSTRAST genome and the *D. virilis* genome (File S1) for sequences matching the SUH-d12 motif, 5′-YGTGRGAAH, in any orientation (Tables 1), 2, and 3). All sites were taken along with the flanking 400 bp regions, unless the distance to the next Su(H) site was less than 800 bp (*i.e.*, <2x flank distance). In cases where the flanking distance was not the full 400 bp due to this site being adjacent to an edge, the flanking distance was allowed to be shorter than 400 bp. All such Su(H) strings were then sequentially named with an lcl # fasta header with lcl numbering beginning with the *D. melanogaster* strings and continuing the same numbering for the *D. virilis* strings. This resulted in unique lcl ID #s for all Su(H) strings regardless of genomic origin (each # beginning with 073 in reference to ″BATCHFETCH 2.0″ run version ″7.3″). Each Su(H) string header also includes information as to the number of Su(H) sites (″S″) in the string and whether the begining and ending flanking sequences were the full 400 bp (″F″ at the beginning, ″f″ at the end) or less (″X″ at the beginning, ″x″ at the end).

**Table 1.**
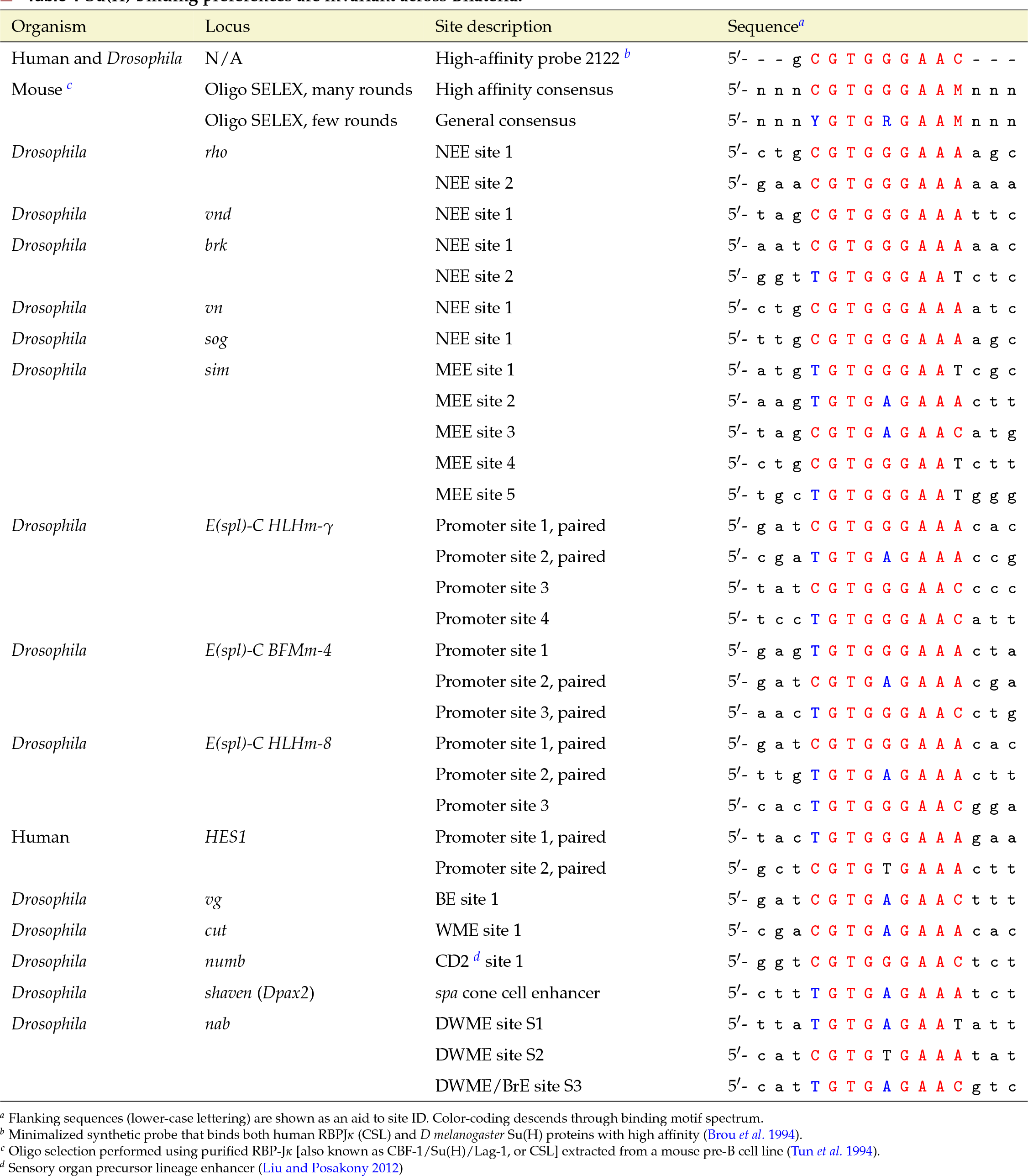
Su(H) binding preferences are invariant across Bilateria.

**Table 2.** Universal Su(H) binding motif spectrum.

| Motif name<sup>a</sup> | Binding affinity range | Motif<sup>b</sup> |
| --- | --- | --- |
|  |  | 5'- 1 2 3 4 5 6 7 8 9 |
| SUH-d2 | high-affinity | 5'- C G T G G G A A M |
| SUH-d8 | ≥ med. aff. | 5'- <u>Y</u> G T G <u>R</u> G A A M |
| SUH-d12<sup>c</sup> | ≥ med. low aff. | 5'- Y G T G R G A A <u>H</u> |
| suh-d18 | > low aff. | 5'- Y G T G <u>D</u> G A A H |
| suh-d144 | > very low aff. | 5'- Y <u>R</u> T G D G <u>W</u> <u>R</u> H |
| Mutated<sup>d</sup> | no binding | 5'- Y c a t D G W R H |

<sup>a</sup> Motif number indicates number of degenerate (d) sequences matching each motif. Higher numbered motifs match a greater number of sequences while continuing to match high-affinity sequences.
<sup>b</sup> IUPAC DNA consensus code used. Underlining indicates increased degeneracy relative to previous motif in spectrum.
<sup>c</sup> This is the motif used in our computational screen.
<sup>d</sup> The 5'-GTG → 5'-CAT mutated sequence is used here to disrupt/knock-down DNA binding by Su(H) as previously shown Tun et al. \(1994\).

**Table 3.** Identification of conserved regulatory belts with Su(H) sites (SUH-RBs).

| Input/Step | Description/Parameter | Output/Value |
| --- | --- | --- |
| Genome assemblies: | *D. melanogaster* BDGP R5/dm3 | 14 contigs |
|  | *D. virilis* r1.3 | 13530 contigs |
| Non-protein-coding filter: | r5.51 FlyBase Genes ∩ CONTRAST | "I-CONTRAST" (24120 blocks) |
| BATCHFETCH 2.0 <sup>a</sup>: | Su(H) (CSL) binding site descriptor | 5'- Y G T G R G A A H |
| | Single site block length (flank length) <sup>b</sup> | 809 bp (Su(H) site ±400 bp) |
| | Max distance between adjacent sites | 799 bp (flank length $-$1 bp) |
|  | Su(H) strings from *D. mel.* I-CONTRAST | 8325 |
|  | Su(H) strings from *D. vir.* | 12058 |
| RBLASTN <sup>c</sup>: | Number of *D. mel.* query hits to *D. vir.* with E-value > E-06 | 1780 belts (*D. mel.*) |
| Unique *D. mel.* SUH-RBs: | Duplicate hits to *D. vir.* removed <sup>d</sup> | 1344 SUH-RBs <sup>e</sup> |
| Multicopy SUH-RBs | Duplicate SUH-RBs (exact duplicate sequences) | 21 SUH-RBs |
| Multicopy transposons | Duplicate sequences matching LINEs or LTRs (RepeatMasker) | 19 SUH-RBs <sup>f</sup> |
| Transposon-overlapping set | SUH-RBs overlapping an annotated transposon by at least 50% | 82 SUH-RBs <sup>g</sup> |

<sup>a</sup> This is an improvement over BATCHFETCH 1.0 (Brittain *et al.* 2014) to better handle multi-site clusters (Materials and Methods).
<sup>b</sup> Some blocks are shorter due to flank edge effects.
<sup>c</sup> Regulatory BLASTN parameter set: penalty -4, reward 5, word size 9, gap open 8, gap extend 6, xdrop gap final 90, best hit overhang 0.25, best hit score edge 0.1 (Brittain *et al.* 2014).
<sup>d</sup> Spot inspection of duplicates indicates that these correspond to LINE and LTR elements.
<sup>e</sup> These are composed of 1157 belts with a single Su(H) site, 158 belts with two sites, and 29 belts with 3–5 clustered sites.
<sup>f</sup> Includes 6/316 mdg1 I gypsy-class, 2/184 Rover-I gypsy-class, 4/1,548 Cr1a LINEs, 5/1,111 I-element LINEs, and 2/350 TART telomere-class LINEs. Denominators indicate number of RepeatMasker database entries for each family (Smit *et al.* 2013). Based on overlap with 27129 LTR (19029) and LINE (8100) entries.
<sup>g</sup> Based on overlap with 81518 entries from the RepeatMasker data set without microsatellite repeats or simple sequence repeats.

Both Su(H) string data sets in fasta format were then used to construct local BLAST data sets for use in a regulatory-specific ″RBLASTN″ parameter set previously described (Brittain *et al.* 2014). An E-value cut-off of E-06 was chosen because this allowed retrieval of all four, non-homologous, canonical NEEs, which are conserved across the genus (NEEs at *rho, vn, brk*, and *vnd*) and corresponds to the amount of conservation associated with a regulatory belt containing approximately 20 binding sites.

**Assigning coordinates:** Bowtie2 was used to map conserved DNA fragments to the *Drosophila melanogaster* r5.51 genome. Only genomic assignments that mapped the entire DNA fragment to the genome with no mismatches were considered as potential locations where the fragments mapped. As some fragments contained identical sequences, they were randomly assigned genomic locations that were subsequently confirmed by hand using the UCSC genome browser BLAT. A BED file based on these coordinates is available in File S2.

**Determining overlapping coordinates:** To determine the overlap between different SUH-RB subsets, we wrote a Python script that compared the genomic coordinates of various data sets used in this study (Table 5 and File S3). Coordinates were considered to overlap if they shared at least one base pair (File S4). We also used this script to determine the total number of unique ChIP-seq peaks between the four data sets [Su(H) 0-8 embryonic ChIP-seq, Su(H) 8-16 embryonic ChIP-seq, Su(H) 16-24 embryonic ChIP-seq, and Su(H) Kc Cells ChIP-seq] (File S4). Peaks that overlapped were counted as one merged peak.

### Molecular cloning

DNA fragments were amplified from genomic DNA extracted from *w*^1118^ flies and cloned into the Xba I site of the pH-Stinger vector (TATA-box containing *hsp70* core promoter driving nuclear eGFP) (Barolo *et al.* 2004) or the EcoR I site of the −42 *eve-lacZ* pCaSpeR vector. Enhancer fragments were sequenced in both directions to confirm identity of clones and the absence of unwanted mutations. Mutations of individual Su(H) sites were created using two-step PCR-mediated stitch mutagenesis to introduce changes as indicated in the text, and sequenced to confirm these mutations. Fragments were amplified using a high-fidelity Taq polymerase (NEB Platinum Taq mix). The following oligonucleotide primer pairs were used to amplify fragments cloned into the pH-Stinger vector: *A2bp1* MlntE: 5′-GGAAAAATCTTAAGCCCTTGC and 5′-ACAGG CGTTATTGTTATGACC; *Delta* WME: 5′-TTCAACGAATGTAAGCGGCC and 5′-TATGCTCCTCTTCAAAGCGC; *esg* 14-16ds (ALE-PLE): 5′-CGCTTATTC AGAACACTTTCCAGG and 5′-ATTCATTGATGCGGAAGATTATAGG; *esg* 16ds (ALE): 5′-GATTTTAGCCCCAAACATTACTTCG and 5′-ATTCATTGATGCGG AAGATTATAGG; *hh* MlntE: 5′-CTTATGTATATTTCCCAAGATTACTTA and 5′-GCACAAAAATAGTAGTAAGC; *knrl* fragment: 5′-TGATTCATGGAAGAGG CCC and 5′-CAAGATTTGCACTGTGATACTCG; and *Notch* WME: 5′-CCA TCCCATGAGATCTTGG and 5′-GATTGGTCGACTTGTGTGG. The following primer pairs were used to amplify fragments cloned into the −42 *eve-lacZ* vector: *E2f1* WPE: 5′-AGAGTTCTTCTCCTCGCTGG and 5′-GCCA TAACCAATACCTTTTGTTCG; and *rst* DicE: 5′-TGGCACGATTACACAGAAAA A and 5′-GGTTTTTGCTCTTCGGATATAA;

### Dissections and tissue staining

For antibody staining, wandering third instar larvae were dissected and fixed with 11.1% formaldehyde in PBS for 30 minutes, followed by 3-4 PBT washes over 30 minutes. Then tissue was then blocked with 1% BSA in PBT for 1 hour. Tissue was incubated with primary antibodies over night, with secondary antibodies for 1.5 hours, and with 17.5 mM DAPI in PBT for 5 minutes. After each incubation, a series of washes was done for 30 minutes. Subsequently imaginal discs were dissected from the remaining tissue and cuticle. This dissection was done on a slide in 80% glycerol, covered with a supported cover slip and imaged with a confocal microscope. The following primary antibodies were used: chicken anti-GFP (1:250) (abcam: ab13070) and mouse anti-β-galactosidase/40-1a (1:12) (developed by R. Sanes, obtained as a supernatant from the Developmental Studies Hybridoma Bank created by the National Institute of Child Health and Human Development of the NIH, and maintained by the University of Iowa Department of Biology). Primary antibodies were detected with Cy2-conjugated goat anti-chicken (1:1000) (abcam:ab6960) or Cy5-conjugated goat anti-mouse (1:1000) (Invitrogen:A10525) secondary antibodies.

For *β*-Gal staining, wandering third instar larvae were dissected and fixed with 1% glutaraldehyde in PBS for 15 minutes, followed by 2 PBT washes 10 minutes each. Dissected tissues were then incubated in an X-gal staining solution (7.2 mM N*a*_2_HP*O*_4_,2.8 mM Na*H*_2_PO_4_,150 mM NaCl, 1 mM MgC*l*_2_, 3 mM *K*_3_[Fe(CN)_6_], 3 mM *K*_4_[Fe(CN)_6_], and 0.21% X-gal) at 37° overnight. Subsequently, imaginal discs were dissected and prepared as described above. The tissue was then imaged using bright field microscopy.

### *Notch* mutant assays

Functional analysis of enhancer constructs in *Notch* mutant background was performed by crossing reporters lines with *N*^1^/*FM7c* flies (Bloomington stock #6873). Larvae with GFP-positive muscle expression indicating the presence of the *FM7c* balancer were compared with larvae with GFP-negative muscle expression.

### Data availability

Transgenic lines carrying enhancer reporter constructs described in this study are available upon request. DNA sequences for novel enhancers shown in figures will be deposited with GenBank along with annotations indicating polymorphisms relative to the reference genome iso-1.

File S1 (FILE-S1-PIPELINE.zip) is a zipped archive of the computational pipeline (scripts and output files) for BATCHFETCH, I-CONTRAST, RBLASTN, and fasta-formatted file of the 1344 SUH-RBs. File S2 (FILE-S2-BED1344.txt) is a BED-formatted file of the 1344 SUH-RBs using coordinates to the Apr. 2006 (BDGP R5/dm3) assembly. File S3 (FILE-S3-overlapscript.txt) is a commented Python script for identifying overlap between different subsets of 1344 SUH-RBs. File S4 (FILE-S4-MASTERLIST.xlsx) is an Excel file containing the master list of information on the 1344 SUH-RBs. File S5 (FILE-S5-SUHRBs-TRANSPOSONS.txt) is a text file of multi-copy SUH-RBs and their transposon families if applicable. File S5 (FILE-S6-FIGS1-knrl.pdf) is a pdf file containing Figure S1 and figure legend and concerns the truncated SUH-RB cloned from the *knrl* intronic region.

## RESULTS

### Identification of Su(H)-binding regulatory belts in *Drosophila*

To identify genome-wide regulatory regions targeted by Su(H) in diverse tissues and developmental stages, we used the neurogenic ectoderm enhancers (NEEs) located at *vnd, rhomboid, vein*, and *brinker* as a training data set (Fig. 1). Su(H) binding sequences (red boxes in Fig. 1) are embedded in a clustered series of ~15-30 peaks of sequence conservation, corresponding to other TFs working with Su(H) in a tissue and stage specific manner. Such series of conservation peaks are frequently well separated from adjacent series, which typically correspond to adjacent *cis*-regulatory modules (CRMs) (*e.g.*, Fig. 1D). In the absence of experimental information identifying whether any such series of non-protein-coding conservation corresponds to a transcriptional enhancer, core promoter, insulator, or other type of CRM, we will refer to the cluster as a (non-protein-coding) regulatory belt.

**Figure 1.**
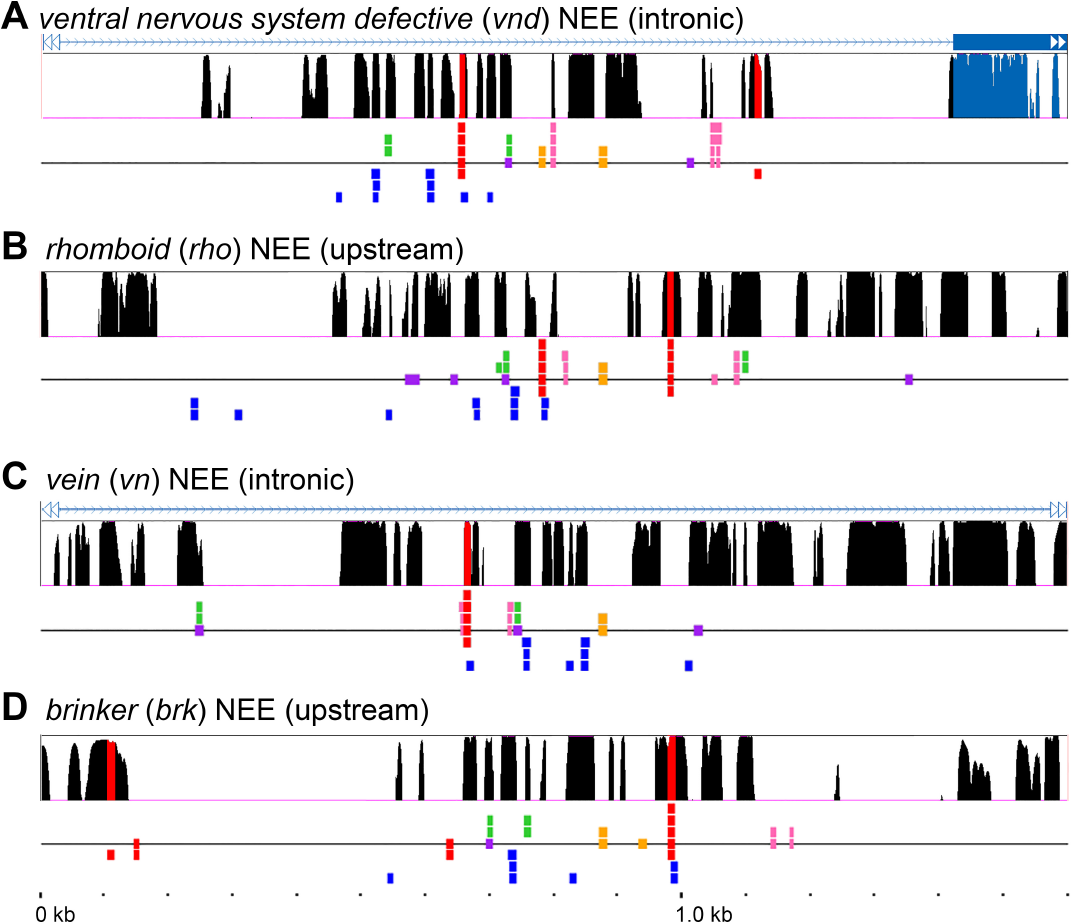
Regulatory belts overlapping the neurogenic ectoderm enhancers (NEEs) provide an example set of non-homologous enhancers using a type of enhancer logic integrating Su(H)/Notch signaling. The NEEs at *vnd* (**A**), *rho* (**B**), *vein* (**C**), and *brk* (**D**) share binding sites for Su(H) (red boxes), as well as Dorsal (blue boxes), Twist:Daughterless (green boxes), Dip3 (orange boxes), Zelda (pink boxes), and Snail (purple boxes). Stacked boxes on higher tracks correspond to more stringent motifs recognizing more specialized and/or higher binding affinity sites. Each panel also shows 12-way insect conservation (black, red, and light blue peaks). Conservation peaks containing Su(H) binding sequences are highlighted in red while peaks overlapping protein-coding sequences are highlighted in light blue. Conserved peaks in the non-protein coding regions correspond in size to TF binding sites and illustrate the typical clustered “regulatory belt” of ~15-30 TF binding sites present seen in conserved enhancers. All panels are the same scale as indicated by ruler (**D**).

The NEE-bearing *vnd* locus illustrates that diverse regulatory belts corresponding to enhancers and promoters are: (*i*) similar in length (800-1000 bp), (*ii*) characterized by a similar number of conservation peaks (15-30 peaks), and (*iii*) well-separated from one another (Fig. 2A). To identify the contextual rules affecting Su(H) binding site usage in enhancers, we used the characteristics of *Drosophila* regulatory belts to identify genus wide RBs containing Su(H) binding sequences. The Su(H) binding motif is ideal for such computational-based comparative genomic screen for at least three reasons. First, the Su(H) gene has generally resisted duplication and diversification in comparison to many other transcription factor families since its evolutionary origination with Metazoa. Thus, vertebrate Su(H), RBPJκ (also CBF1 or CSL for CBF1/Su(H)/Lag-1), has the same biochemical DNA binding preference as *Drosophila* Su(H), which demonstrates that the Su(H) target sequence and binding preferences are highly constrained (Tun *et al.* 1994). Second, it is not a member of a large eukaryotic transcription factor superfamily as are the C_2_H_2_ zinc finger-containing, homeodomain-containing, and bHLH families, each of which represent TFs with often continuously overlapping binding preferences. To estimate the extent to which Su(H) binding sequences are unique for Su(H), we ran different Su(H) binding sequences through the JASPAR Insecta TF binding profiles. We find that no other TF binding profiles match any of the Su(H) binding sequence with a score > 85%. Third, several known Notch-target enhancers and their functional Su(H) binding sites have been extensively characterized in *Drosophila* (Table 1).

**Figure 2.**
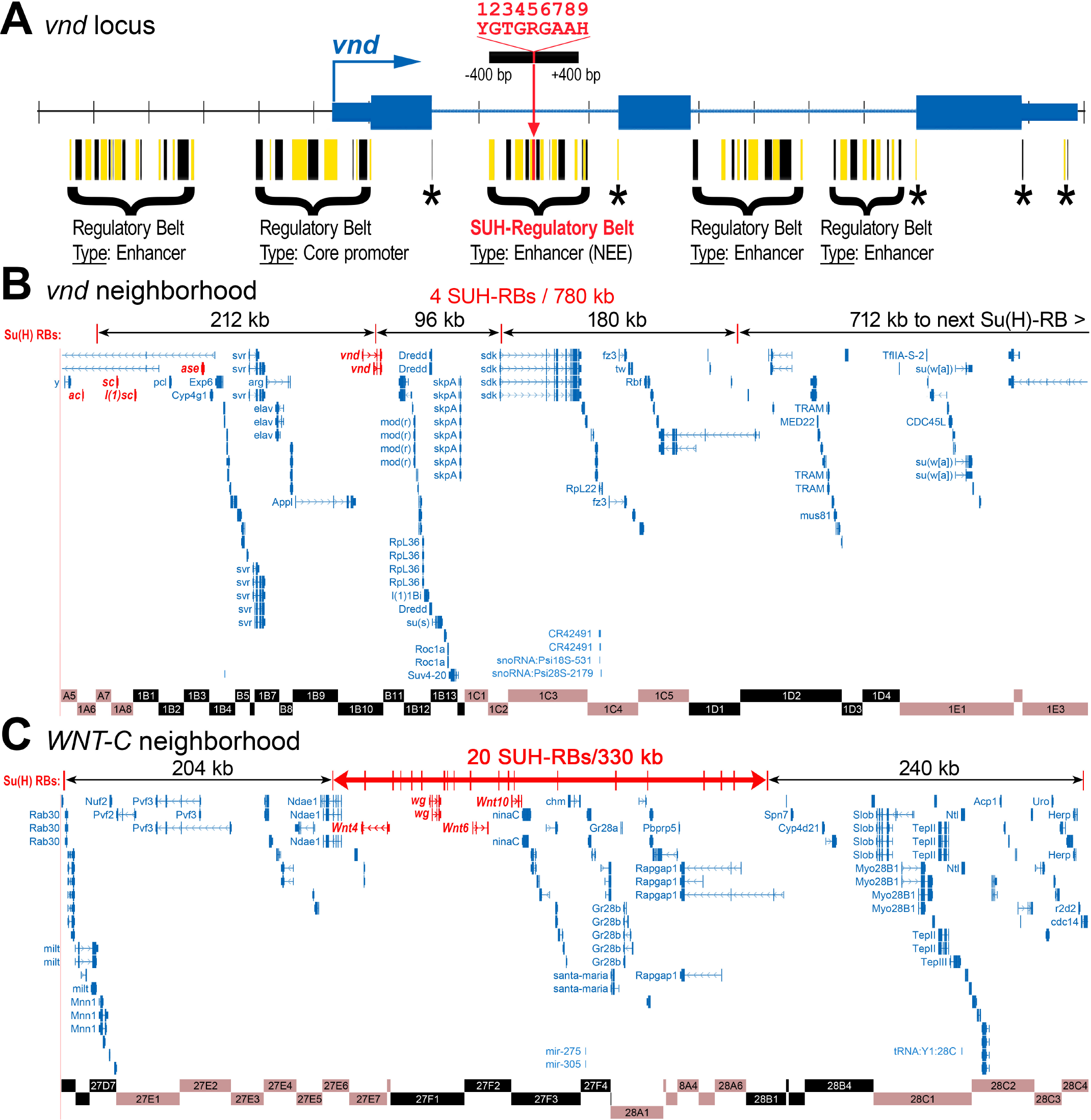
Characteristic regulatory belts with Su(H) binding sequences are unevenly distributed across the genome. (**A**) The conserved nonprotein coding regions of the *vnd* locus (*D. melanogaster* shown) demonstrates the typical dimensions of regulatory belts, which correspond to transcriptional enhancers and core promoters (clustered peaks of conservation in a 0.8-1.0 kb window). Peaks are shown in alternating red and black as a visual aid. Other types of regulatory sequences, such as intronic splice sites or 3′ UTR sites (asterisks), are not embedded in regulatory belts, which have substantially higher amounts of sequence conservation. In this study, we identify all regulatory belts containing one or more canonical Su(H) binding sequences (±400 bp), such as the neurogenic ectoderm enhancer (NEE) present in the first intron of *vnd* (Table 1). (**B**) The nearest Su(H)-containing regulatory belts (RBs) in the *vnd* locus (red lines) are located 100-200 kilobases or more away from the *vnd* NEE as shown in this 780 kb window. Chromosomal bands are shown below the gene tracks. (**C**) In contrast, other genomic regions, such as this 780 kb window containing the four *WNT* genes of *Drosophila (Wnt4, wg, Wnt6*, and *Wnt10*), can have extremely dense super-clusters of SUH-RBs. In this particular case, a 330 kb window around the Wnt-encoding genes contains 20 SUH-RBs. This unexpected high density of SUH-RBs at the *WNT* gene cluster is comparable to the high density of SUH-RBs at the E(*spl*)-complex (not shown). The *WNT* super-cluster of SUH-RBs is well separated from the next adjacent SUH-RBs at distances similar to the *vnd* genomic neighborhood (B). In total, we identified 1344 SUH-RBs present in the euchromatic portion of the *D. melanogaster* genome.

We used known in vivo Su(H) binding sites and biochemical data from human, mouse, and fly (Table 1) to determine an optimal Su(H) search sequence within the context of a binding motif spectrum (Table 2). This ″SUH-d12″ motif matches the Su(H) sites in canonical Notch target enhancers (Table 1). The first position in the Su(H) binding site can be a pyrimidine, but is a ′C′ in the high-affinity site. Furthermore, in the characteristic paired inverted sites present in promoters from the *Enhancer of split* complex *(E(spl))* or the homologous vertebrate *HES* promoters, one site always begins with a ′C′ and the other with a ′T′. This suggests that this first position is a functional ″inflection″ of Su(H) site grammar. The fifth position can be a purine, but is a ′G′ in the high-affinity site. The ninth position can be an ′A′ or a ′C′ in medium to high-affinity sites, however sometimes it is a ′T′ in combination with other positions with high-affinity nucleotides. Just as important, the ninth position is rarely ever a ′G′ (Table 1). In the NEEs, there is typically a single high-affinity Su(H) binding site with few neighboring Su(H) sites, if any (Erives and Levine 2004). Thus, this NEE Su(H) binding site features a ′C′ in position 1, a ′G′ in position 5, and typically an ′A′ in position 9 (Crocker *et al.* 2010). These observations from the NEEs, coupled with the Su(H) site inflections in paired inverted sites (Bailey and Posakony 1995) and the biochemical selection experiments (Tun *et al.* 1994), suggest that Su(H) binding sequences are differentially constrained in different contexts. We therefore used a medium-affinity Su(H) motif 5′-YGTGRGAAH (see SUH-d12 in Table 2) to search for all genomic windows with a matching sequence in the relatively large *D. virilis* genome and in a non-protein-coding version of the smaller *D. melanogaster* genome (see Materials and Methods). To ensure, Su(H) binding sequences were retrieved as part of full-length enhancers, these sequences were trimmed +/- 400 bp from either side of the Su(H) binding sequence.

The use of a filtered *D. melanogaster* genome allowed us to lower our expectation value for a reciprocal RBLASTN between the two Su(H) data sets until all five dorsal-Twi-Su(H) targeted neurogenic ectoderm enhancers (NEEs) were identifiable (see Materials and Methods). The five NEEs are functionally conserved across the genus at the *vnd, brk, sog, rho*, and *vein* loci and typically feature a single high-affinity Su(H) site (Erives and Levine 2004; Crocker *et al.* 2008, 2010; Crocker and Erives 2013; Brittain *et al.* 2014). Nonetheless, most of these enhancers have experienced binding site turnover so as to make detection unfeasible in a genome by genome nucleotide BLAST query. The *vein* NEE has experienced the most turnover between the two lineages, which represent the two *Drosophila* sub-genera, without losing functionality (Crocker *et al.* 2008). In the RBLASTN module of our pipeline, the *D. melanogaster vein* NEE is identified as being conserved in *D. virilis* at an E-value of 5e-07, which is the highest E-value of all the NEEs (Table 3). We thus set our RBLAST threshold to be E-06, and identified 1344 evolutionarily-conserved regulatory blocks with Su(H) binding sites, of which 1157 are single site blocks and 187 are blocks containing between 2-5 sites for a collective total of 1575 SUH-d12 sites (Table 3).

We also determined that over 98.4% of the 1344 SUH-RBs are single copy sequences consistent with unique regulatory regions (Table 5). A spot check of the 436 filtered SUH-RBs in *D. melanogaster* that had multiple hits in *D. virilis* SUH-RBs indicated that these were repeats (Table 3). We then investigated whether any of the remaining 21 multi-copy SUH-RBs (duplicate within *D. melanogaster*) overlap any of the tens of thousands of known transposable elements (Bartolomé *et al.* 2002; Smit *et al.* 2013). We report that 19 of these SUH-RBs correspond to multi-copy transposable elements (RepeatMasker data set, see Methods) in the SUH-RB data set (Table 5). These correspond to six mdg11 gypsy-class LTRs, two Rover-I gypsy-class LTRs, four Cr1a LINEs, five I-element LINEs, and two TART telomere-class LINEs. Moreover, as each of these classes of transposable elements exist in hundreds of copies (Bartolomé *et al.* 2002), retro-elements with Su(H) binding sequences are not representative of entire families of LINEs or LTR-type retro-elements. Nonetheless, it is possible that some of these SUH-RBs might belong to active elements of a transposon family or pseudogenized retro-elements undergoing sequence drift and/or co-option into host regulatory functions (Feschotte 2008; Sundaram *et al.* 2014).

### Su(H) binding sites are constrained to be medium affinity sites

To investigate the extent of non-local binding site constraints and possible functional ″inflections″ in Su(H) binding sequences, we analyzed all 1575 SUH-d12 sites in the 1344 SUH-RBs (Fig. 3A). For example, we find and report on two specific SUH-d12 sequence patterns in which only the fifth position purine is allowed to ″wobble″ (Fig. 3B). Of the sites matching the sequence motif 5′-<u>T</u>GTG<u>R</u>GAA<u>A</u>, over 60% feature a fifth position G. In contrast, of the sites matching the sequence motif 5′-<u>C</u>GTG<u>R</u>GAA<u>C</u>, the frequency of a fifth position G drops down to about 44%. Importantly, the first position C and fifth G choices both lead to higher affinity binding sequences and are seen together in biochemical SELEX experiments (Tun *et al.* 1994). Thus, these results suggest that Su(H) binding sequences are under selection for avoiding the most high affinity possible sequences. It also further suggests that selection at certain TF:site wobble positions can compensate for other positions.

**Figure 3.**
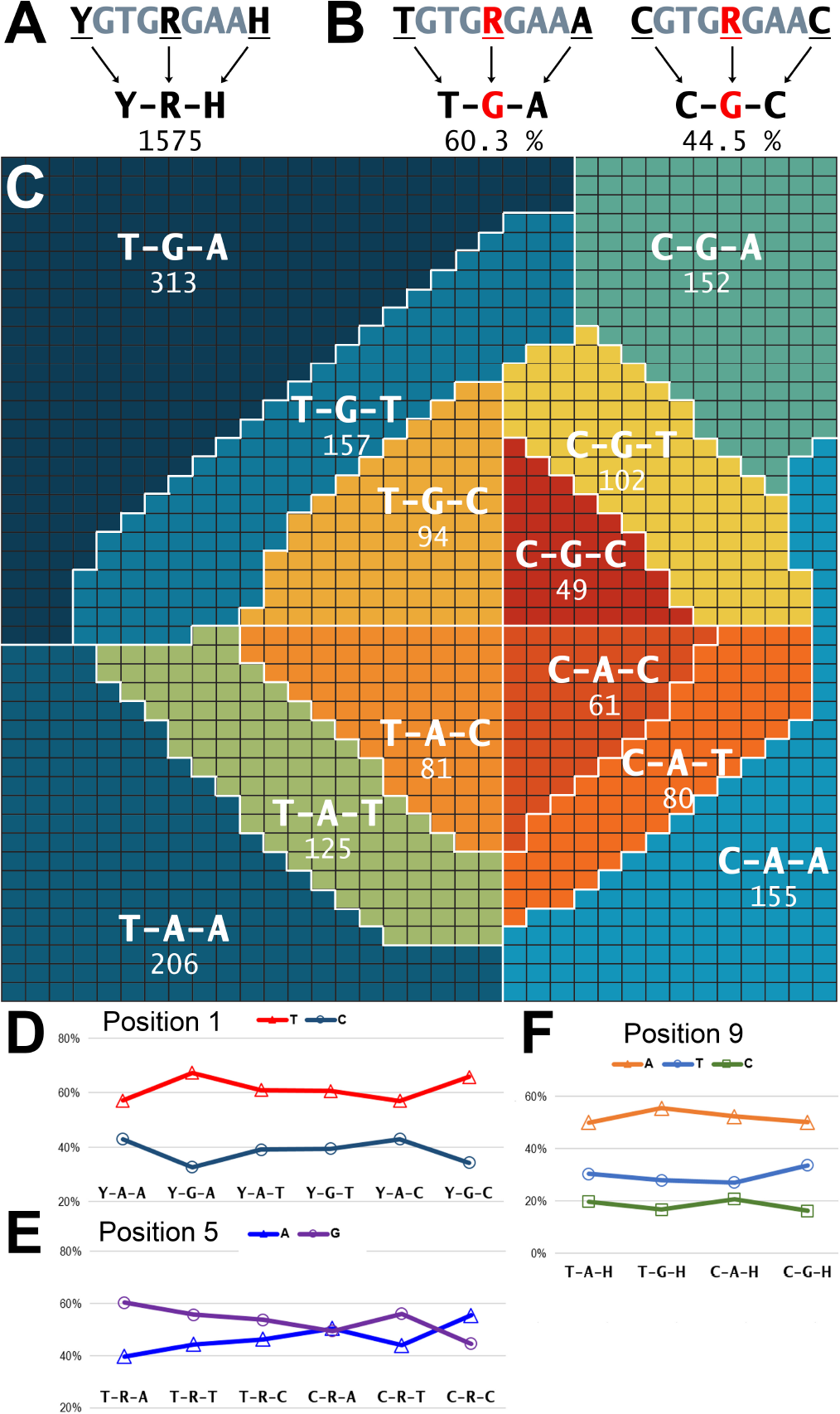
Genomic dimensions of Su(H)-binding site inflection. (**A**) The SUH motif used in the SUH-RB screen features three wobble positions (“Y-R-H”), which results in 12-fold degeneracy. The 1344 SUH-RBs harbor 1575 SUH sites. (**B**) Shown are two sub-SUH sequences restricted to fifth position wobble (puRines “A” and “G”). In the “T-R-A” pattern, R = “G” at 60.3 %, and corresponds to the most common SUH sequence. However, in the “C-R-C” pattern, R = “G” only 44.5% of the time, suggesting that this sequence constrains this wobble position. See Results section for significance. (**C**) This grid is a representation of all 1575 SUH sequences present in the 1344 SUH-RBs. Each unit square represents a single site and is color-coded by the specific SUH sequence (1/12 possible sequences). The heat map colors the most numerous (dark blue) to the least numerous (red) sequence classes, which are arranged so that position 1 is divided into “T” sequences on the left-hand-side and “C” sequences on the right-hand-side; position 5 is divided into “G” sequences on top and “A” sequences on the bottom; and position 9 is divided into concentric rings with “A” sequences in the outer ring, “T” sequences in the middle ring, and “C” sequences in the inner diamond. (**D-F**) These graphs show the percentage of sequences matching the indicated pattern given the presence of a particular letter at positions 1 (**D**), 5 (**E**), or 9 (**F**). Positions 5 and 1 appear to be the most constrained relative to the nucleotides present in adjacent wobble positions.

To get a global picture of the 1575 Su(H) binding sequences found within the 1344 SUH-RBs, we plotted all the sites as a function of their type and frequency (Fig. 3C). This overview demonstrates the preference for medium affinity sequences at functional sites as follows. In the first position of the binding site, the less optimal T is featured in about 62% (976/1575) rather than the higher-affinity C. In the ninth position of the binding site, the less optimal T is featured 1.6 x as much as the higher-affinity C. However, this is not the case for the ninth position A, which is the most frequent. A constraint for GC-context, a possible alternate explanation, does not ring true as the fifth position G is favored in 55% of the 1575 sites.

To investigate the extent of wobble-choice among the 1575 Su(H) binding sites, we also graphed the percentage of sequences featuring a certain letter at a wobble position relative to all sequences matching a sequence pattern (Fig. 3D-F). Non-local constraints are greatest at position five, then position one, and last position nine because these positions feature differences between the maximum and minimum frequencies of 15.8%, 10.3%, and 4.4-6.6%, respectively.

### SUH-RBs are unevenly distributed across the genome

The 1344 Su(H)-type regulatory belts (SUH-RBs) are unevenly distributed across the genome. For example, a 780 kb window around the vnd locus shows that in addition to identifying the intronic NEE, only three other Su(H) site containing RBs are identified (Fig. 2B). These are located from 100-700 kb away from each other.

In stark contrast, some chromosomal regions contain dense-clusters of SUH-RBs. Some of these dense SUH-RB regions are expected, such as the *E(spl)* gene complex, which includes many canonical Notch targets using inverted paired Su(H) sites (see Table 1). The region encompassing *E(spl)*-C contains 21 SUH-RBs in a 280 kb window, thus averaging 7.5 SUH-RBs/100 kb.

Other genomic regions that are not known to be as intimately connected to Notch signaling also have comparable densities of SUH-RBs. For example, a 330 kb window encompassing the *Drosophila WNT* genes (*Wnt4, wg, Wnt6*, and *Wnt10*) contains a total of 20 SUH-RBs, averaging 6.1 SUH-RBs/100kb (Fig. 2C). In *Drosophila*, Notch signaling functions upstream of *wg* in dorsal/ventral (D/V) wing margin establishment (Diaz-Benjumea and Cohen 1995; Neumann and Cohen 1996). Notch signaling also regulates *WNT* genes in other systems from hydra (cnidarians) to vertebrates (Naylor and Jones 2009; Münder *et al.* 2013). Thus, our finding of multiple SUH-RBs over the entire fly *WNT* cluster may indicate that Notch regulates Wnts in diverse developmental contexts.

In another example of dense SUH-RB clustering, a 144 kb window encompassing the related genes *knirps (kni)* and *knirps-like (knrl)* contains 9 SUH-RBs, thus averaging 6.2 SUH-RBs/100 kb (Fig. S1A). In this example, the next closest SUH-RBs are found 238 kb and 89 kb away from the ends of this SUH-RB cluster. Similar to the *WNT* gene cluster, this inter-cluster spacing suggests that the SUH-RBs are the result of general Notch involvement in gene regulatory networks (GRNs) involving Knirps-family factors.

Last, we find that the *Delta* locus, which encodes one of two membrane-bound ligands for the Notch receptor (Knust *et al.* 1987; Vässin *et al.* 1987), harbors 9 SUH-RBs in 99 kb, thus averaging 9.1 SUH-RBs/100 kb. In comparison, the *Notch* locus has only a single SUH-RB, suggesting that Notch-Delta signaling may be extensively modulated by regulation of the genes encoding Notch ligands. We therefore also looked at the *Serrate (Ser)* locus, which encodes the second Notch receptor ligand in *Drosophila* (Fleming *et al.* 1990). We find that the *Ser* locus harbors 6 SUH-RBs in 32 kb. Thus, the two genes encoding Notch receptor ligands, *Delta* and *Ser,* together have a SUH-RB density of 11.5 SUH-RBs/100 kb (15 SUH-RBs/131 kb). All together, these results suggest that the genomic distributions of SUH-RBs identify the key developmental genes constituting Notch-regulated GRNs.

### Su(H)-dependent enhancers at *Notch* and *Delta*

Our first use of the 1344 SUH-RBs was to identify the SUH-RB with the densest cluster of Smad Dpp/BMP-effector, Apterous, and Zelda binding sites (Stroebele and Erives 2016). This led to the identification of the *nab* dorsal wing margin enhancer (DWME) Stroebele and Erives (2016). A single SUH-RB at *Notch* and one of the *Delta* SUH-RBs also contained a similar set of sites as the *nab* DWME. We therefore assayed these two SUH-RBs to see if they drove similar expression patterns in wing imaginal discs.

We cloned a 573 bp SUH-RB fragment containing one Su(H) binding from the *Notch* locus (Fig. 4A). This fragment drives expression in the pouch of the developing wing imaginal disc (Fig. 4B). This expression is limited to the D/V and anterior/posterior (A/P) compartment boundaries and a quadrant-type pattern. Along the anterior to posterior (A-P) axis there is a gap in reporter expression corresponding to peak Dpp expression (Fig. 4B). However, this *Notch* enhancer is not affected by a *Notch^1^* mutant background (Fig. 4 compare control background in C to mutant background in D). Nonetheless, a mutated *Notch* enhancer in which the single Su(H) binding site is changed to a non-binding sequence (5′-CGTGAGAAT → 5′-CcatAGAAT) drives very weak expression (Fig. 4 compare E to F and G). Thus, the single SUH-RB at *Notch* is a complex wing margin enhancer (WME) requiring an intact Su(H) binding site to drive robust expression throughout the wing pouch.

**Figure 4.**
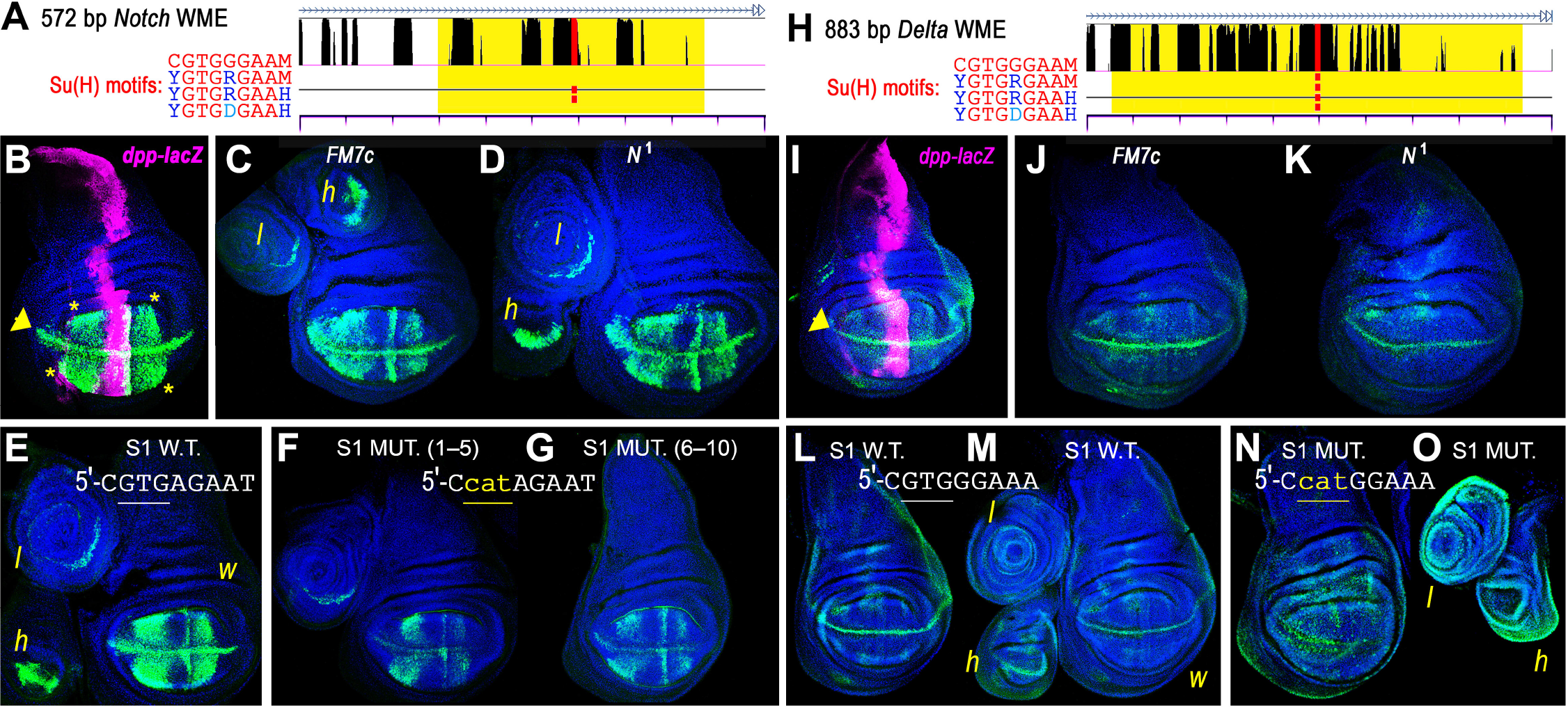
Conserved SUH-RBs at *Notch* and *Delta* are Su(H)-dependent wing margin enhancers (WMEs). We searched the 1344 SUH-RB sequences for matches to binding motifs for Mad:Medea (BMP effectors), Zelda (pioneer factor expressed in embryos and imaginal discs), and Apterous (dorsal compartment wing disc selector) and found regulatory belts in the *Notch* (**A-G**) and *Delta* (**H-O**) loci. (**A**) The 572 bp cloned SUH-RB (yellow highlight) from the second intron of *Notch* contains a single medium affinity Su(H) binding sequence. Matches to the four Su(H) binding site motifs shown are indicated on separate tracks with the third motif, which was used in the computational pipeline, shown on the black line. (**B**) The cloned *Notch* fragment is a wing imaginal disc enhancer driving GFP expression in a wing pouch quadrant (yellow asterisks) and along the D/V wing margin (yellow arrowhead). The disc was dissected from third instar larvae and double-stained for *Notch* WME-driven GFP (green) and β-gal (magenta) expressed by a *dpp-lacZ* reporter. The *Notch* WME appears to read-out Notch and Dpp/BMP signals independently in wing discs. It also drives expression in a broader D/V margin-like stripe in haltere discs (h) and a partial ring stripe in leg discs (l) as seen in adjacent panels. (**C, D**) Shown is activity from the *Notch* WME in either an *FM7c* (**C**) or *N*^1^ mutant background (**D**) suggesting that wild-type Notch receptor is not absolutely required for margin expression. (**E-G**) Shown is imaginal disc activity from the *Notch* WME (**E**) or a mutated *Notch* WME with an inactivated Su(H) binding site (**F, G**), which drives precipitously attenuated expression as seen in these representative discs from independent pools of lines 1-5 (**F**) and lines 6-10 (**G**). (**H-O**) Equivalent panels for the *Delta* regulatory belt (yellow highlight), which we cloned and found to be another WME. (**H**) Shown is a region of the first intron of *Delta* with the cloned fragment highlighted in the yellow box and the Su(H) sites shown in red. It also appears to be a D/V margin enhancer in haltere discs (**M**). Unlike the *Notch* WME, the *Delta* WME drives a more diffuse expression pattern throughout the wing pouch. The D/V wing compartment margin expression driven by the *Delta* WME appears to be slightly more affected by a mutant *N*^1^ background (**K**) but is still not absolutely dependent on wild type Notch signaling. (**N,O**) Mutation of the Su(H) site in the *Delta* WME results in the loss of D/V margin expression in both wing and haltere discs and a gain of ectopic expression in wing discs (**N**), and leg and haltere discs (**O**).

We cloned an 883 bp SUH-RB fragment containing one Su(H) binding site from the *Delta* locus (Fig. 4H). This fragment drives robust expression along the entire D/V compartment boundary (presumptive wing margin region) and more diffusely throughout the wing pouch (Fig. 4I). This *Delta* WME expression pattern weakens only slightly in a *Notch* mutant background, as seen when we compared 10 discs for each of the control and mutant backgrounds (compare representative discs in Fig. 4J and K). Nonetheless, a mutated enhancer in which the Su(H) binding site is changed to a non-binding sequence (5′-CGTGGGAAA → 5′-CcatGGAAA) no longer drives expression along the margin (Fig. 4 compare L and M to N). In addition, this mutated *Delta* enhancer now drives ectopic expression throughout the leg and haltere imaginal discs (Fig. 4 compare M to O). The mutated *Delta* enhancer also drives some weak ectopic expression outside the wing pouch in wing imaginal discs (Fig. 4 compare L and M to N). Thus, this particular SUH-RB at *Delta* is a canonical WME requiring an intact Su(H) binding site to drive expression at the margin. Unlike the *Notch* WME, the *Delta* WME requires its Su(H) binding site for repression outside of the presumptive wing margin and in other imaginal discs. See Discussion for further details.

### Identification of novel SUH-RB enhancers from different stages

In addition to the *Notch* and *Delta* WMEs, we cloned and tested 11 SUH-RBs and found that 10 of these drove distinct tissue-specific activities, which we show in this study and companion studies (*nab* DWME (Stroebele and Erives 2016) and 9 novel enhancers in the upper and bottom halves of Table 4, respectively). In a companion study on the regulation of *nab* developmental gene (Stroebele and Erives 2016), we found that a SUH-RB containing two Su(H) sites in the first intron on *nab* corresponds to a wing/haltere imaginal disc enhancer (DWME) and a larval brain enhancer (BrE) (Table 4). The *nab* DWME is licensed by wing disc selectors to integrate Notch and Dpp/BMP signaling pathways and drive expression in the dorsal wing margin. The *nab* DWME activity is also attenuated by the Su(H) site in the adjacent BrE and was one of several identified inter-enhancer silencing activities observed in the *nab* intronic enhancer complex.

**Table 4.** SUH-RBs corresponding to known Notch-target enhancers (top) and novel enhancers identified in this study (bottom).

| Locus | lcl_073x | # sites<sup>a</sup> | Enhancer/activity | References<sup>b</sup> |
| --- | --- | --- | --- | --- |
| *vnd* | 460 | 1 | Neurogenic ectoderm enhancer (NEE) | (Erives and Levine 2004; Brittain *et al.* 2014) |
| *brk* | 822 | 3 | Neurogenic ectoderm enhancer (NEE) | (Markstein *et al.* 2002; Erives and Levine 2004; Crocker *et al.* 2008) |
| *cut* | 838 | 1 | Wing margin enhancer (WME) | (Jack *et al.* 1991; Guss *et al.* 2001) |
| *sog* | 1286 | 1 | Neurogenic ectoderm enhancer (NEE) | (Crocker *et al.* 2008) |
| *numb* | 1659 | 1 | Sensory organ precursor lineage (CD2) | (Rebeiz *et al.* 2011) |
| *Su(H)* | 2852 | 4 | Auto-regulatory socket enhancer (ASE) | (Barolo *et al.* 2000) |
| *vg* | 3961 | 1 | Wing margin enhancer (boundary element, BE) | (Williams *et al.* 1994; Kim *et al.* 1996) |
| *rho* | 4767 | 2 | Neurogenic ectoderm enhancer (NEE) | (Ip *et al.* 1992; Erives and Levine 2004; Markstein *et al.* 2004; Crocker *et al.* 2010) |
| *nab* | 5382 | 2 | 764 bp dorsal wing margin enhancer DWME<sup>c</sup><br>288 bp larval brain enhancer (BrE) | (Stroebele and Erives 2016)<br>(Stroebele and Erives 2016) |
| *vn* | 5477 | 1 | Neurogenic ectoderm enhancer (NEE) | (Erives and Levine 2004; Brittain *et al.* 2014) |
| *sim* | 7238 | 5 | Mesoectoderm enhancer (MEE) | (Kasai *et al.* 1992; Cowden and Levine 2002) |
| *hh* | 7792 | 1 | Lymph gland enhancer (*hh*-f4F) | (Tokusumi *et al.* 2010) |
| *dysf* | 7964 | 1 | Leg imaginal disc enhancer (GMR_13D07<sup>d</sup>) | (Jory *et al.* 2012; Cordoba and Estella 2014) |
| *E(spl)mδ-HLH* | 7948 | 2<sup>e</sup> | Unknown | (Bailey and Posakony 1995; Nellesen *et al.* 1999) |
| *E(spl)mγ-HLH* | 7949 | 2* | Embryonic ventral neuroectoderm, imaginal sensory organ (non-neuronal) | (Bailey and Posakony 1995; Nellesen *et al.* 1999) |
| *E(spl)mγ/mβ* | 7950 | 2* | Unknown | (Bailey and Posakony 1995) |
| *E(spl)mβ-HLH* | 7952 | 2 | Embryonic ventral neuroectoderm, imaginal proneural | (Nellesen *et al.* 1999) |
| *E(spl)mα-BFM* | 7954 | 5 | Imaginal SOPs | (Nellesen *et al.* 1999; Castro *et al.* 2005) |
| *E(spl)m3-HLH* | 7956 | 2* | Unknown | (Nellesen *et al.* 1999) |
| *E(spl)m4-HLH* | 7957 | 3* | Imaginal proneural | (Bailey and Posakony 1995; Nellesen *et al.* 1999) |
| *E(spl)m7-HLH* | 7960 | 4* | Imaginal proneural | (Singson *et al.* 1994; Nellesen *et al.* 1999) |
| *E(spl)m8-HLH* | 7961 | 3* | Imaginal proneural, wing margin | (Kramatschek and Campos-Ortega 1994; Bailey and Posakony 1995; Nellesen *et al.* 1999; Cave *et al.* 2005; Janody and Treisman 2011) |
| *rst* | 138 | 1 | 782 bp “DicE” = dice pips enhancer (DicE): punctate embryonic expression | This study (Fig. 5A) |
| *Notch* | 590 | 1 | 578 bp wing margin enhancer (WME): wing/haltere/leg discs | This study (Fig. 4A–G) |
| *esg* | 2862 | 2 | 1352 bp larval epidermal enhancer (LEpE) | Manuscript “a” planned |
|  | 2865 | 1 | 3098 bp posterior larval (sensory) enhancer (PLE)<sup>f</sup> | This study (Fig. 5D) |
|  | 2866 | 1 | 1144 bp anterior larval (sensory) enhancer (ALE)<sup>g</sup> | This study (Fig. 5C) |
| *rho* | 4765 | 1 | 1217 bp tracheal placode enhancer (TPE) | Manuscript “b” in preparation |
| *A2bp1* | 5727 | 1 | 638 bp larval/adult midgut intestinal enhancer (mIntE) | This study (Fig. 5E) |
| *Delta* | 7577 | 1 | 884 bp WME: wing/haltere/leg discs | This study (Fig. 4H–O) |
| *E2f1* | 7720 | 1 | 582 bp wing pouch enhancer (WPE): pouch region of wing and haltere imaginal discs | This study (Fig. 5B) |
| *hh* | 7793 | 1 | 1430 bp adult midgut intestinal enhancer (MIntE) compartment (*hh*-D) | This study (Fig. 5F) |
|  | 7794 | 1 | 547 bp larval imaginal disc enhancer (*hh*-E) | Manuscript “c” in preparation |

<sup>a</sup> Only SUH-d12 sites are counted here. Asterisk (\*) indicates paired inverted SUH-d12 sites separated by 15 bp.
<sup>b</sup> References pertain to one of the following: (i) identification of enhancer, (ii) evidence supporting direct regulation by Su(H), and/or (iii) evidence supporting membership in a mechanistically-equivalent set of enhancers regulated by Su(H) (inverted pair of Su(H) sites in *E(spl)*-C promoters or NEE-type site organization).
<sup>c</sup> The “*nab*-A” fragment is 764 bp in the reference genome, and 762 bp in the *w*<sup>1118</sup> genome.
<sup>d</sup> This 3.7 kb *Janelia* driver encompasses the SUH-RB and the minimalized *Dys640* enhancer, which are 642 bp away from each other.
<sup>e</sup> Paired inverted sites separated by 13 bp instead of 15 bp.
<sup>f</sup> Tested as 14ds–16ds fragment, which has ventral posterior larval cells + ALE activity
<sup>g</sup> Tested as 16ds and 14–16ds fragments, where “#ds” = # kb downstream of *esg*

We cloned a 782 bp SUH-RB fragment from the second intron of *roughest (rst)* (Table 4). This cloned fragment drives robust *lacZ* reporter expression in a dynamic series of changing ″pips″ in late embryonic series (Fig. 5A). We refer to this enhancer as the *rst* dice pips enhancer (DicE).

**Figure 5.**
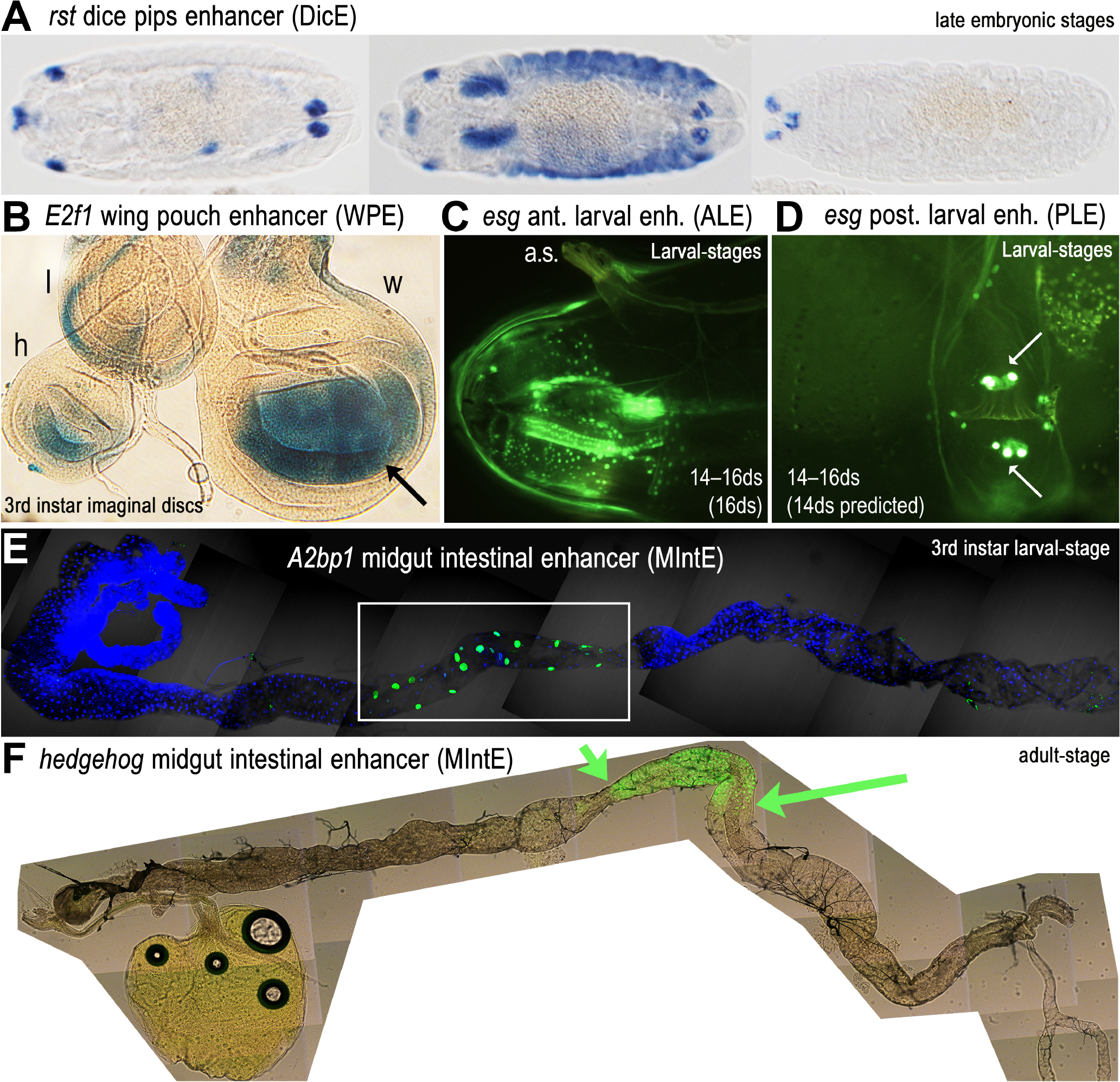
The 1344 SUH-RBs are enriched in many novel enhancers active in different tissues and developmental stages. (**A**) Shown is an anti-sense *lacZ* in situ showing expression driven by an intronic enhancer from the *rst* locus. This “dice pips” enhancer (DicE) drives expression in a dynamic punctate pattern in late embryonic stages. Three representative embryos are shown. (**B**) An intronic enhancer from the *E2f1* locus drives imaginal disc expression, including strong expression in the wing (w) and haltere (h) pouches of imaginal discs (arrow points to wing pouch) as shown by enzymatic β-gal staining. Expression in leg imaginal discs occurs as a ring (l), and in the antennal discs and larval brain (not shown). (**C, D**) Shown are images of live larvae expressing GFP (green) driven by a 3.1 kb fragment that encompasses two adjacent SUH-RBs, which are located 14 and 16 kb downstream (“14-16ds”) of *escargot (esg)*. (**C**) One enhancer activity, which is also seen with a smaller 1.1 kb “16ds” fragment (not shown), drives expression in what appears to be anterior sensory neurons throughout the mouth parts and pharynx. Larval image is a lateral view with anterior to the left and dorsal on top as shown by the extended anterior spiracle (a.s.). (**D**) A second enhancer activity, which is not seen with the 16ds fragment and is therefore attributed to the 14ds regulatory belt, drives expression in a smaller number of possible posterior sensory neurons along the ventral midline. Arrows point to two rows of three neuronal cells. Additional smaller cells can be seen just anterior and posterior to the bigger cells. Faint traces of expression can also be seen in what appears to be neuronal processes. (**E**) Shown is a dissected larval intestinal tract (anterior to the left, posterior to the right) with GFP expression driven by an *A2bp1* enhancer. The tissue is also stained with DAPI (blue). Strong GFP expression is prominent in the large flat cell compartment of the midgut (boxed area). (**F**) Shown is a dissected adult intestinal tract (anterior to the left, posterior to the right) with live GFP expression driven by a *hh* enhancer. Strong GFP expression is prominent in a posterior midgut sub-compartment defined by abrupt anterior (small green arrow) and posterior (large green arrow) boundaries.

We cloned a 682 bp SUH-RB fragment from the first intron of *E2F1* (Table 4). This cloned fragment drives robust *lacZ* reporter expression in the pouch regions of the wing and haltere imaginal discs of third instar larvae, as well as expression in the leg imaginal discs (Fig. 5 B, β-gal X-gal stain).

We cloned a 3098 bp SUH-RB fragment containing two adjacent SUH-RBs located 14 and 16 kb downstream of *escargot (esg)* locus (″14-16ds″, Table 4). This fragment drives expression in third instar larval cells in the anterior mouth and pharyngeal region (Fig. 5 C) and in two rows of three neuron-like cells in the ventral posterior region (see arrows in Fig. 5 D). To determine whether these were separate enhancer activities corresponding to the two adjacent SUH-RBs, we also cloned a smaller 1144 bp fragment of the 3098 bp fragment containing the SUH-RB located 16 kb downstream (″ ′16ds″, Table 4). This 16ds fragment drives the anterior larval expression seen with the 3.1 kb 14-16ds enhancer, but does not drive the posterior larval expression (data not shown). This suggests that the posterior larval enhancer (PLE) corresponds to the 14 kb downstream (″14ds″) SUH-RB. These two *esg* larval enhancers drive anterior and posterior regional expression patterns similar to the late embryonic expression patterns (Whiteley *et al.* 1992).

We cloned a 638 bp SUH-RB fragment from the second intron of *Ataxin-2 binding protein 1 (A2bp1)* (Table 4). This fragment drives expression in the large flat cells of the larval midgut (Fig. 5 E). This *A2bp1* midgut intestinal enhancer (MIntE) also drives expression in the adult midgut (data not shown).

We cloned a 1.4 kb SUH-RB fragment from the *hedgehog* (hh) locus (Table 4). This fragment drives expression in the posterior midgut of the adult intestines (Fig. 5 E). Like the *A2bp1* MIntE, the *hh* MIntE drives expression in a very specific sub-compartment consistent with findings of a highly-regionalized midgut (Marianes and Spradling 2013). Both Notch and Hedgehog signaling are known to be important for intestinal stem cell populations and intestinal regeneration (Marianes and Spradling 2013; Tian *et al.* 2015).

Only a single SUH-RB fragment, a 630 bp cloned DNA from the *knrl* locus, did not drive noticeable reporter activity in larval or adult stages. However, in retrospect this fragment corresponds to a truncated regulatory belt, suggesting that critical elements for activity were not included (Fig. S1).

To determine how many known Notch-target enhancers are present in the 1344 SUH-RBs, we also collated 29 previously characterized CRMs regulated by Notch signaling (Table 4). We find that a majority of these (72.4%, 21/29) are among the 1344 SUH-RBs, suggesting that most known Notch-target enhancers are conserved across the genus (Table 5). In the Discussion section, we review details concerning the 8 known Notch-target enhancers not found in the 1344 SUH-RBs.

**Table 5.**
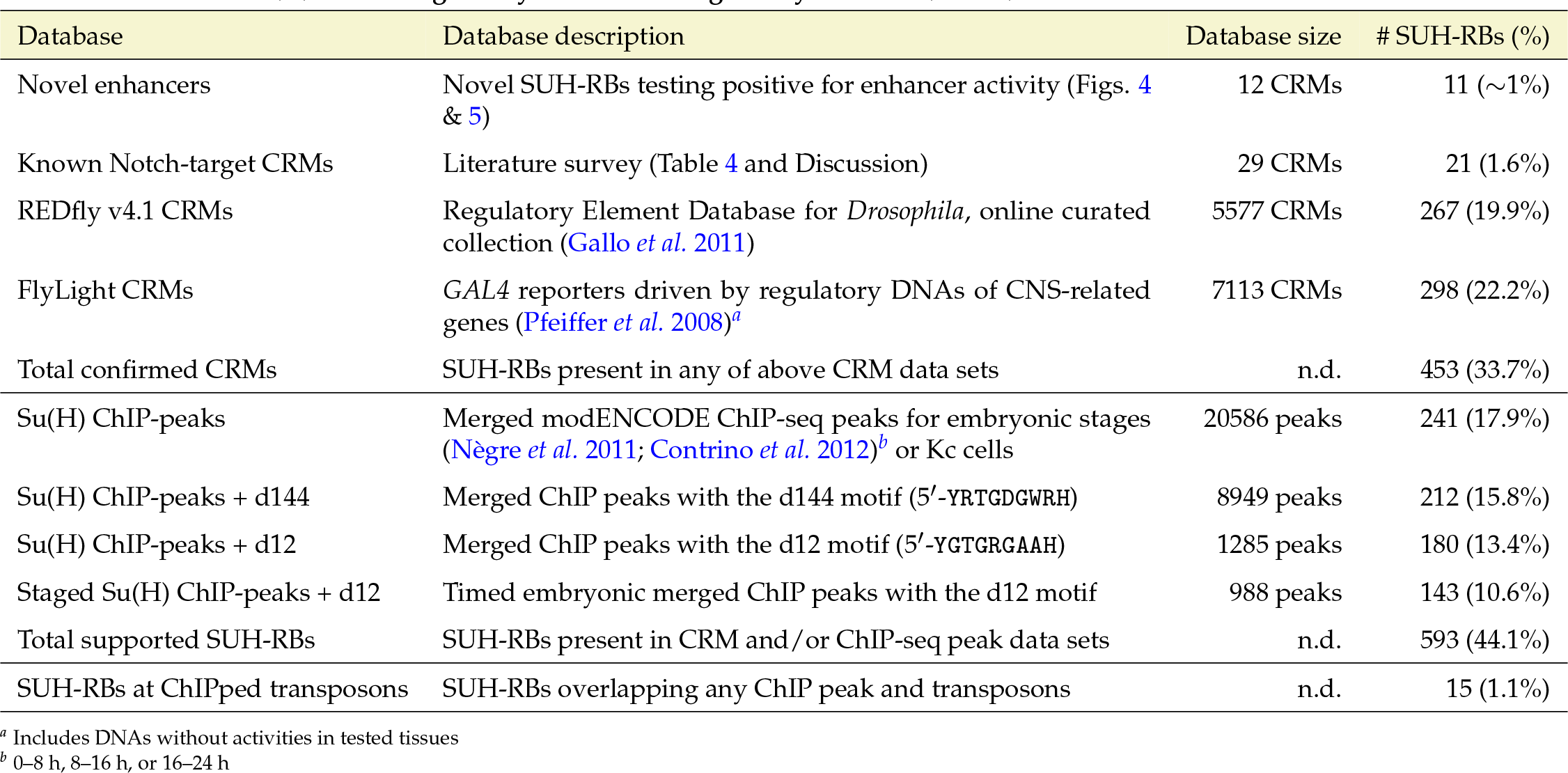
The 1344 Su(H)-related regulatory belts are cis-regulatory modules (CRMs).

### SUH-RBs overlap many known *cis*-regulatory modules

To determine the extent to which the SUH-RBs correspond to known enhancers, we compared them to two enhancer databases: REDfly (Gallo *et al.* 2011) and FlyLight (Pfeiffer *et al.* 2008) (Table 5). The REDfly data set contains curated *Drosophila* enhancers from the literature and annotates whether a fragment is the smallest known active enhancer fragment (*i.e.*, ″minimalized″ CRM). (Gallo *et al.* 2011). This meta-data allows us to avoid over-counting matches due to annotated series of nested enhancer fragments with identical activities. We find that about 20% of SUH-RBs (267/1344) overlap with at least one minimalized enhancer from REDfly (Table 5). However, only 20 of these SUH-RBs correspond to enhancers with curated experimental evidence supporting direct Notch regulation.

The REDfly dataset of *Drosophila* CRMs includes some large-scale enhancer data sets, such as the Vienna TILEs (Kvon *et al.* 2014), for a total of almost 5,600 CRMs. However, REDfly is unlikely to be comprehensive as many more CRMs remain to be characterized. To get an independent measure of SUH-RB/enhancer overlap, we compared the 1344 SUH-RBs to a second large data set not incorporated into REDfly: the FlyLight enhancer-GAL4 reporters, which are constructed from loci relevant to the nervous system (Pfeiffer *et al.* 2008). We find that 22% of SUH-RBs (298/1344) overlap with at least one regulatory region from the FlyLight data set. Only a minority (8.7%) of the SUH-RBs (117/1344) are present in both the REDfly and FlyLight data sets. These comparisons suggest that Su(H)-dependent regulation of enhancers is more extensive than currently known.

Altogether, 33.7% of the SUH-RBs (453/1344) correspond to previously identified enhancer activities or enhancers identified here (Table 5). Furthermore, given that only 1 in 12 cloned enhancers was not found to drive reporter expression, our results suggest that the overwhelming majority of the 1344 SUH-RBs are bona fide enhancers.

### SUH-RBs delineate true from false positive ChIP-seq peaks

A comprehensive inventory identifying all genomic enhancer DNAs for all tissues and developmental stages using reporter assays is not available. Thus, we considered the ways in which chromatin binding data for Su(H) could further inform the nature of the 1344 SUH-RBs. Given the importance of Notch signaling in a discrete number of cells throughout all of development, ChIP-seq data from any one stage could be insufficient to identify all enhancer-related Su(H) binding sites, particularly if binding at these enhancers occurs only in the presence of tissue-specific factors. Therefore, stage-specific ChIP-seq data for Su(H) could tell us the proportion of the 1344 SUH-RBs that correspond to enhancers for that stage.

We compared the SUH-RBs to biochemical Su(H) ChIP-seq binding data for three consecutive embryonic windows (spanning 0-24 h) and embryo-derived Kc cells (Nègre *et al.* 2011; Contrino *et al.* 2012). We find that ~18% (241/1344) of SUH-RBs overlap genomic intervals defined by a merging of all ChIP peaks (Table 5). This suggests that ~82% of the SUH-RBs correspond to enhancers active in stages outside of embryogenesis. Based on our initial reporter assays of the SUH-RBs, this would be consistent with a large proportion of enhancers active in larval imaginal discs and tissues maintained through active stem-cell populations (larval and adult intestines).

ChIP-seq data for Su(H) could also tell us the proportion of ChIP-seq peaks that correspond to “false-positive” peaks that are not located in regulatory belts with any type of Su(H) binding sequence. These ″false-positive″ peaks could be due to (*i*) random associations (artifact); (*ii*) the presence of spurious binding sequences, which likely would be located outside of regulatory belts; or (*iii*) indirect associations resulting from distal looping interactions with Su(H)-dependent enhancers. For example, neighboring Su(H)-targeted CRMs could interact in enhancer-complex silencing hubs as previously suggested (Schaaf *et al.* 2009, 2013; Stroebele and Erives 2016). In this case, Su(H) could bring together distal Su(H)-dependent CRMs (Fig. 6A, left hand-side model). In conjunction with tissue-specific factors (green triangles), Su(H) and the NICD co-activator could free a CRM to allow tissue/stage-specific gene activation (Fig. 6A, right hand-side model). In this case, ChIP-seq data for one stage could help identify Su(H)-dependent CRMs active in other stages, particularly in regions characterized by dense SUH-RBs. Alternatively, enhancer/silencer-hubs could be created by Su(H) binding to only some of the enhancers (Fig. 6 B). In this second case, Su(H) ChIP-seq signals are likely to include CRMs not targeted by Su(H). We thus analyzed ChIP-seq peaks according to those peaks overlapping SUH-RBs, those peaks missing an SUH-d12 sequence but still possessing at least one suh-d144 sequence, and false-positive peaks, which do not overlap SUH-RBs nor contain any suh-d144 sequences ((Fig. 6C).

**Figure 6.**
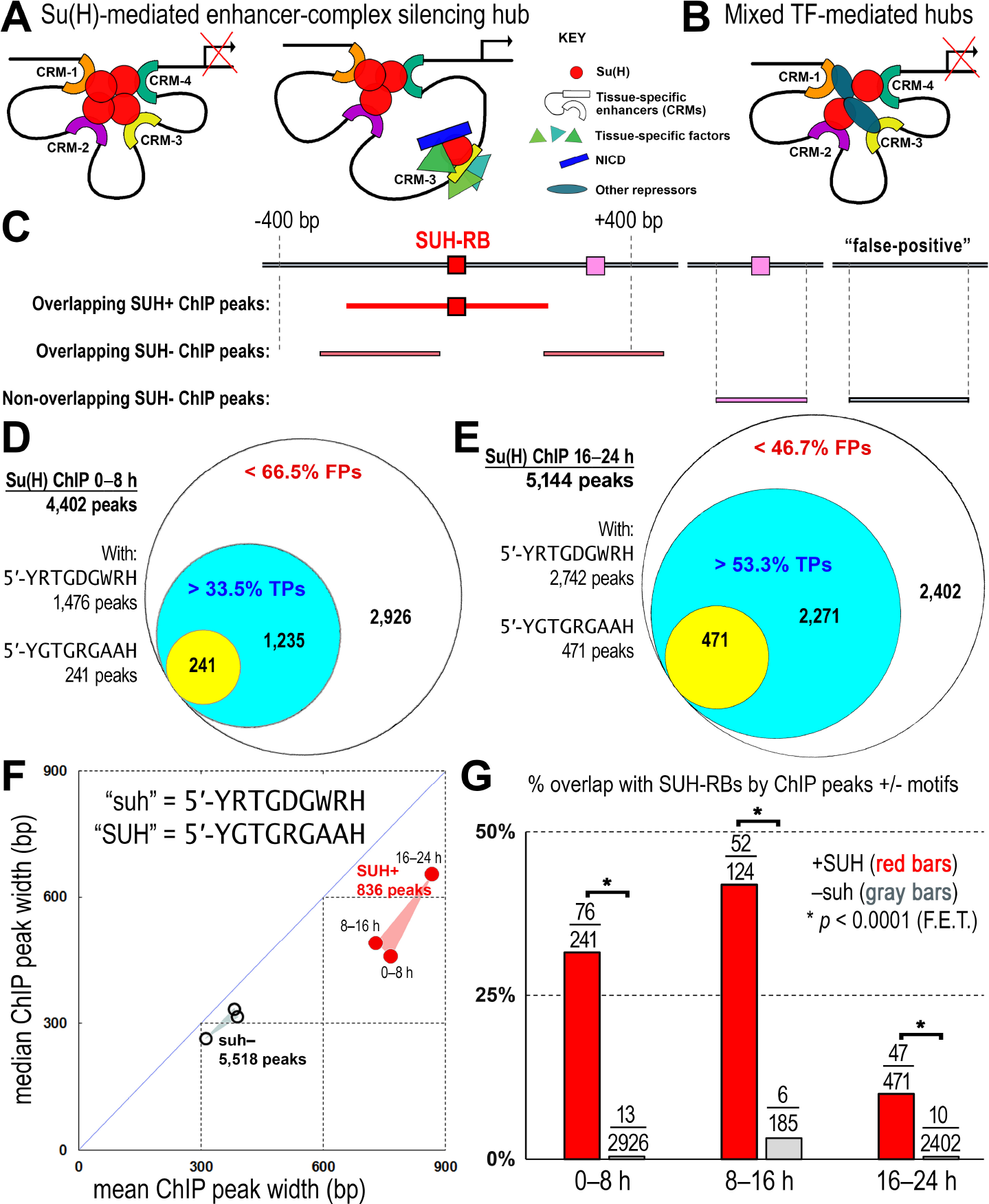
The 1344 Su(H) regulatory belts (RBs) separate true positive (TP) ChIP-seq peaks from false-positive (FP) peaks. (**A**) Various scenarios should be considered in evaluating the overlap of SUH-RBs with tissue and stage specific ChIP-seq data for Su(H). Given the importance of Su(H) and Notch signaling in a discrete number of cells throughout all of development, it is possible that ChIP-seq data from any one stage will be insufficient to identify all enhancer-related Su(H) binding sites, particularly if such binding occurs only in the presence of tissue-specific factors. Alternatively, previous studies indicate that neighboring Su(H)-targeted c/s-regulatory modules (CRMs) may interact in an enhancer-complex silencing hub regardless of the tissue and stage specificity of the CRMs (left hand-side cartoon with CRMs brought together by Su(H) proteins indicated by the red circles) (Stroebele and Erives 2016; Schaaf *et al.* 2013). In conjunction with tissue-specific factors (green triangles), Su(H) (red circle) and the Notch intracellular domain (NICD) co-activator (blue rectangle) could free a CRM to allow tissue/stage-specific gene activation (right hand-side cartoon). Thus, the extent to which Su(H)-targeted enhancers are present in such hubs could determine the extent to which ChIP-seq signals from a single tissue/stage identifies Su(H)-targeted enhancers active in other tissues and stages. (**B**) Illustrated is a mixed TF model in which enhancer/silencer-hubs are created in part by Su(H) binding to only a subset of the enhancers. In this case, Su(H) ChIP-seq signals are likely to include CRMs not targeted by Su(H). (**C**) Shown is a diagram of how we are defining “false-positive” ChIP-seq peaks, which would lack either both high and low affinity Su(H) binding sequences (red and pink boxes, respectively) and not overlap with any of the SUH-RBs. (**D,E**) We find that only 33.5% and 53.3% of Su(H) ChIP peak signals from 0-8 h embryos (**D**) and 16-24 h embryos (**E**), respectively, contain low-(cyan) to high-(yellow) affinity Su(H) binding sites. This suggests lower bounds on the number of “true-positive” Su(H)-binding sites (TPs) and upper bounds on the number of false-positive Su(H)-binding sites (FPs). (**F**) ChIP-seq peaks containing >medium-affinity Su(H) binding sequences (“SUH”-d12 motif) are much longer than Su(H) “peaks” that are missing even a low-affinity Su(H) binding sequence (suh-d144 motif) as the former have much longer ChIP peak means and widths. This is true from three different, non-overlapping embryonic stages (0-8 h, 8-16 h, and 16-24 h). Because SUH-RBs are 800 base pairs or longer in length, these results suggest that false positive ChIP peaks, which have much shorter peak widths than TP peaks, are not the result of adjacent Su(H) binding sites lying just outside of the FP window of sequence but overlapping with the RB window. (*G*) A comparison of the number of ChIP peaks overlapping any of the 1344 Su(H) RBs shows that TP ChIP peaks (+SUH) are much more conserved than FP ChIP peaks (-suh). A Fisher’s Exact T-test (*) confirms the significance of each comparison at each time point (*p* < 0.0001).

To determine the extent of false-positive peaks, we analyzed the sequences of Su(H) ChIP-peak intervals for the early (0-8 h) and late (16-24 h) embryonic stages (Fig. 6D and E). We find that only 33.5% (1235/4402) of peaks from 0-8 h embryos (Fig. 6C) and 53.3% (2271/5144) of peaks 16-24 h embryos (Fig. 6D) contain Su(H) binding sequences of low or greater affinity (suh-d144 motif). ChIP-seq peaks for the intermediate stage (8-16 h) show similar results as the other stages with 53.3% containing at least one suh-d144 sequences (272/510). These results suggest that we could estimate meaningful bounds on the number of false-positive Su(H)-binding sites (FPs). To corroborate this possibility further, we compared ChIP-seq peaks containing medium-affinity Su(H) binding sequences (″SUH+″ peaks containing the SUH-d12 motif in Fig. 6E) to false-positive peaks that are missing even a low-affinity Su(H) binding sequence (″suh-″ lacking the suh-d144 motif in Fig. 6F and Table 2). We find that the SUH+ peaks have much longer ChIP peak mean widths than the suh-peaks for all three embryonic stages (Fig. 6F). This additional result strongly suggests that the presence of an Su(H) binding sequence is sufficient to cut ChIP-seq data into two meaningful classes. Furthermore, because SUH-RBs are 800 base pairs or longer in length, FP ChIP peaks are not the result of adjacent Su(H) binding sites lying just outside of the FP ChIP peak interval (*e.g.*, Fig. 6C).

We thus proceeded to determine the extent of ChIP-peak overlap with SUH-RBs according to the presence of canonical Su(H) binding sequences (SUH+) or absence of possible Su(H) binding sequences (suh-). We find that true positive ChIP peaks (+SUH) overlap SUH-RBs much more frequently than FP ChIP peaks (-suh) (Fig. 6G). This difference is significant as determined by Fisher′s Exact Test (*p* < 0.0001 in all cases).

Almost 90% of the SUH-RBs that overlap any Su(H) ChIP peaks (212/241) contain a sequence matching the suh-d144 motif (Table 5). Most of these are retained if the more stringent SUH-d12 motif is used as ~75% of SUH-RBs overlapping ChIP peaks (180/241) contain a SUH-d12 sequence. This d12 motif, which we used in our screen, is also likely to have a lower false-positive rate of enhancer detection than the d144 motif for the following reason. Only 1.2% of the ChIP-peak data set overlaps with the 1344 SUH-RBs (241/20586, see Table 5). This percent overlap doubles to 2.4% (212/8949) when only peak intervals with sequences matching the d144 motif are considered. However, this percent increases to 14.0% (180/1285) when only sequences matching the d12 motif are considered.

Last, we checked the triple intersection of the 1344 SUH-RBs, all Su(H) ChIP peaks, and transposable elements. We find that only 15 SUH-RBs (1.1%) intersect both data sets, suggesting that the 1344 SUH-RBs are predominantly single-copy regulatory regions unrelated to transposable elements (Table 5).

## DISCUSSION

In this study, we identified 1344 enhancer-like regulatory belts containing canonical binding sites for Su(H) and conserved across the *Drosophila* genus. Of these 1344 SUH-RBs, ~34% overlap bona fide enhancers most of which are not known to be direct targets of Notch/Su(H) regulation (Table 5). Including the SUH-RBs with embryonic and Kc ChIP-seq peaks, this number increases by an additional 10% to ~44% of SUH-RBs that have experimental support (Table 5). In terms of the 29 known Notch-target enhancers, the 1344 SUH-RBs overlap 21 of them (72.4%) (Table 4). The 8/29 known Notch-target enhancers neither overlap the 1344 SUH-RBs nor lie within 1 kb of them. These 8 pertain to the following enhancers: the *broad (br)* early enhancer (brE) (Fuchs *et al.* 2012; Jia *et al.* 2014), a *gcm* glial enhancer (Jones *et al.* 2004), a *hedgehog* posterior wing margin enhancer (Pérez *et al.* 2011), a *salm* wing enhancer (Guss *et al.* 2001), a *Ser* wing enhancer (Yan *et al.* 2004), the *Sox15* sensory organ socket cell enhancer (Miller *et al.* 2009), the *sparkling (spa)* cone cell enhancer of *shaven (sv)* (Fu *et al.* 1998; Swanson *et al.* 2010), and the *vg* quadrant enhancer (QE) (Guss *et al.* 2001). However, many of these loci have other non-overlapping SUH-RBs elsewhere in the locus.

Analysis of some of the unidentified Notch-target enhancers illustrates different reasons for their absence in the set of 1344 SUH-RBs. One example is the *sparkling* eye cone cell enhancer harbored at the *sv* locus, which encodes D-Pax2 (Fu *et al.* 1998; Swanson *et al.* 2010). Inspection of the *spa* enhancer fragments shows that the 655 bp enhancer fragment contains one d-12 site and two d-144 sites (see Table 2), while a minimalized 362 bp fragment contains only the two d-144 sites. Furthermore, we find that *spa* is unrecognizable as a conserved regulatory belt in the distantly related *D. virilis* genome. This is consistent with previous studies indicating a rapid evolutionary turnover of binding sites at this enhancer in lineages more closely related to *D. melanogaster* (Swanson *et al.* 2011). Thus *spa* is not identified as an SUH-RB in this study for two independent reasons: a lack of conservation and the use of non-SUH-d12 low-affinity sites in *D. melanogaster*. Similarly, the brE features Su(H) binding sites in *D. melanogaster* that do not match the SUH-d12 motif (Fuchs *et al.* 2012; Jia *et al.* 2014). Thus, the downstream gene battery directly regulated by Notch and Su(H) in any given fly lineage may actually be greater than the set of 1344 enhancers identified here. This would be due to some enhancers not being conserved across the genus and/or some enhancers containing low-affinity, non-SUH-d12 sites in either *D. melanogaster* or *D. virilis*.

In addition to identifying the Notch/Su(H)-dependent *nab* DWME (Stroebele and Erives 2016), we also identified Su(H)-dependent WMEs at *Notch* and *Delta* among the 1344 SUH-RBs (Fig. 4). However, the activities of these two enhancers in the *Notch*^1^ mutant background were not as impacted as the *nab* DWME (Stroebele and Erives 2016). Interpretation of these WME-driven expression patterns in the *Notch*^1^ versus *FM7c* balancer backgrounds is complicated by two aspects of these backgrounds. First, the reason for the semi-dominant hypomorphic nature of the *Notch*1 mutant is unknown and is complicated by several additional mutations accrued since its discovery (Lehmann *et al.* 1983; Dietrich and Campos-Ortega 1984). Second, the *FM7c* balancer encodes an expanded polyglutamine tract Q_15_HQ_17_ at Notch relative to wild-type Notch (Q_13_HQ_17_), which may be functionally important (Rice *et al.* 2015). We also identified 9 additional distinct enhancers from the SUH-RBs, 6 of which we have shown here (Fig. 5), and 3 of which we will report in separate, more detailed manuscripts in preparation (Table 4). Further work will be needed to identify the specific roles played by Su(H) and Notch signaling for each of these enhancers.

While most Notch/Su(H)-target enhancers were identified as belonging to the 1344 SUH-RBs, we were surprised to find overlap with many known enhancers not previously found to be targeted by Su(H) and/or Notch signaling. This suggests that Notch-signaling cross-talk with other signaling pathways and/or Su(H)-mediated regulation may be generally featured across more developmental enhancers than has been previously appreciated.

The vast majority of SUH-RBs that have been found to be enhancers suggests there is a negligible false-positive rate of enhancer detection. We found that only one of the 12 novel SUH-RBs that we cloned did not have any GFP reporter expression in larval or adult stages (see *knrl* in Table 4). This “larval FP” regulatory belt is located in an intron of *knirps-like (knrl)* and is one of nine SUH-RBs clustered in a 144 kb window encompassing *knrl* and *knirps (kni)* (Fig. S1A). Two possible reasons could explain the reason for our not detecting activity. First, this cloned regulatory belt could be active in other stages or cells that we did not investigate, or in a single cell or small group of cells that are easily missed in the larval or adult stages, which we did investigate. Second, the 630 bp cloned fragment is centered on a single SUH-d12 site that sits on the edge of a distinct regulatory belt of conservation (Fig. S1B). Thus, this cloned fragment could be missing critical elements contained in the remaining part of the regulatory belt, including some lower affinity Su(H)-binding sequences (red boxes in Fig. S1B). In conclusion, our results on the genomic dimensions of SUH-RBs suggest that these are all likely to be enhancers for various developmental stages and/or adult stem-cell populations.

## ACKNOWLEDGMENTS

This work was supported in part by an NSF CAREER award to AE to study morphogen gradient enhancer readouts (IOS:1239673), NIH T-32 bioinformatics training grant support to ES, two Evelyn Hart Watson research fellowships to ES (in 2014 and 2015), and an Iowa Biosciences Academy (IBA) fellowship to TF. We thank Crystal Maki for assistance in cloning the *E2f1* enhancer. We thank Daniel Eberl for consultation with the expression of *esg* enhancers.

